# Metabolomic Profiling and Characterization of a Novel 3D Culture System for Studying Chondrocyte Mechanotransduction

**DOI:** 10.1101/2024.06.10.598340

**Authors:** Priyanka P. Brahmachary, Ayten E. Erdogan, Erik P. Myers, Ronald K. June

## Abstract

**Background/Objective:** Articular chondrocytes synthesize and maintain the avascular and aneural articular cartilage. *In vivo* these cells are surrounded by a 3D pericellular matrix (PCM) containing predominantly collagen VI. The PCM protects chondrocytes and facilitates mechanotransduction. PCM stiffness is critical in transmitting biomechanical signals to chondrocytes. Various culture systems with different hydrogels are used to encapsulate chondrocytes for 3D culture, but many lack either the PCM or the *in vivo* stiffness of the cartilage matrix. This study aimed at establishing a culture system to investigate a) if chondrocytes cultured in alginate will develop a PCM and b) study mechanotransduction via metabolic changes induced in 3D agarose-embedded chondrocytes upon mechanical stimulation.

**Methods:** We cultured primary human and bovine chondrocytes in monolayers or as alginate encapsulated cells in media containing sodium L-ascorbate. PCM expression was analyzed by immunofluorescence and western blots. We further characterized the response of chondrocytes embedded in physiologically stiff agarose to dynamic compression through metabolomic profiling.

**Results:** We found that primary human and bovine chondrocytes, when cultured in alginate beads with addition of sodium L-ascorbate for 7 days, had a pronounced PCM, retained their phenotype, and synthesized both collagens VI and II. This novel culture system enables alginate-encapsulated chondrocytes to develop a robust PCM thereby creating a model system to study mechanotransduction in the presence of an endogenous PCM. We also observed distinct compression-induced changes in metabolomic profiles between the monolayer-agarose and alginate-released agarose-embedded chondrocytes indicating physiological changes in cell metabolism.

**Conclusion/Significance:** These data show that 3D preculture of chondrocytes in alginate before encapsulation in physiologically stiff agarose leads to pronounced development of pericellular matrix that is sustained in the presence of ascorbate. This model can be useful in studying the mechanism by which chondrocytes respond to cyclical compression and other types of loading simulating *in vivo* physiological conditions.

## Introduction

Osteoarthritis (OA), the most common form of arthritis, is a chronic degenerative joint condition affecting more than 37% of people over the age of 60. It is a leading cause of pain and disability resulting in considerable productivity loss and healthcare expenditure [1]. Pathological changes in OA include progressive degradation and loss of articular cartilage, thickening of the subchondral bone, inflammation of the synovium, degeneration of ligaments and hypertrophy of the joint capsule [2]. What was once considered a simple “wear and tear” condition, is now known to be a disease of the whole joint with low-grade inflammation and other biological mechanisms playing a role in disease progression [3, 4].

The articular cartilage that lines diarthrodial joints is primarily composed of the extracellular matrix (ECM) and the pericellular matrix (PCM). Specialized cells of cartilage, chondrocytes, occupy 3-5% of tissue volume in adult human cartilage. They synthesize and maintain the avascular and aneural cartilage and are surrounded by a 3D pericellular matrix that facilitates mechanotransduction. Chondrocytes also maintain a functional extracellular matrix by synthesis and assembly of the collagen fibers, proteoglycans, and other matrix molecules [5, 6]. During regular joint loading *in vivo*, cartilage is subjected to mechanical stresses that can exceed 10 megapascals [7].

However, chondrocytes in healthy cartilage are protected from these mechanical loads by the PCM. In contrast to OA chondrocytes have a softer PCM. Chondrocytes can sense and respond to mechanical stimuli, but the mechanisms by which this mechanotransduction occurs are not fully understood [8, 9]. Understanding the basic processes involved in chondrocyte response to mechanical loads is likely to provide valuable insights for the treatment of osteoarthritis.

Various culture systems with different hydrogels are used to encapsulate chondrocytes and study mechanotransduction, but many lack either the PCM or the *in vivo* stiffness of the cartilage matrix [10–12]. These hydrogels have stiffness values ranging from 5-10 kPa [13]. PCM stiffness in healthy adult chondrocytes is between 25-200 kPa, which significantly decreases in OA [14, 15]. Physiological stiffness is important since the hydrogel directly transmits mechanical information to chondrocytes, driving the biological responses via cellular mechanotransduction. Cyclical compression can upregulate transcription of ECM genes depending on the compression amplitude and frequency [16, 17]. Similarly, mechanical forces also play a role in gene expression of aggrecan and several other matrix proteins [18, 19].

Stoddart *et al* reported that chondrocytes cultured in alginate beads will form a pericellular matrix and display a phenotype similar to *in vivo* conditions [20]. The alginate gel can be effectively removed using chelating agents to facilitate the release of the resulting chondrons (*i.e.* chondrocytes with PCM) making it a useful system to study extracted chondrocytes with an endogenously synthesized PCM. Therefore, the primary objective of this study was to determine if alginate-encapsulated chondrocytes could develop a PCM and subsequently be encapsulated in physiologically stiff agarose. This culture system aims both to promote PCM formation and to serve as a functional model for studying chondrocyte mechanotransduction. The resulting metabolic responses of chondrocytes subjected to mechanical stimuli may lead to a better understanding of the molecular mechanisms underlying mechanotransduction with significant implications for cartilage tissue engineering.

These data indicate that primary human and bovine chondrocytes, when cultured in alginate beads with the addition of sodium L-ascorbate, develop a pronounced PCM, synthesize collagens VI and II, and retain their phenotype. In contrast, chondrocytes cultured as monolayers did not form a distinct PCM. Our findings show that 3D culture of chondrocytes in alginate using complete media supplemented with ascorbate leads to increased production of matrix proteins, collagens VI and II. We observed that both primary osteoarthritic human and normal bovine chondrocytes, when cultured as a monolayer in ascorbate showed a pronounced pattern of secreted collagens VI and II resembling a PCM. Encapsulating the chondrocytes in alginate beads, and culturing in ascorbate resulted in chondrons that synthesized substantially more collagen VI and II near the perimeter of the cell. In monolayers cultured with ascorbate, Collagen VI and II expression appears punctuate, forming discreet foci around the chondrocytes. This punctate pattern suggests that collagen secretion is localized and forms small, discontinuous regions of pericellular matrix (PCM) around individual cells. The punctate appearance may reflect the limitations of a 2D environment, where chondrocytes are spread out and lack a 3D structure to support uniform collagen deposition. While the presence of ascorbate enhances the secretion of collagen, the resulting PCM-like structure in monolayer culture is less robust and lacks the continuous chondron-like organization seen in alginate cultures, suggesting that monolayer culture might not fully replicate the biomechanical and spatial cues of a native cartilage environment.

Encapsulating chondrocytes in alginate beads and culturing with ascorbate results in a more uniform and intense collagen deposition near the cell periphery resembling native chondrons. Collagens VI and II are synthesized in greater amounts compared to monolayer cultures, likely due to the 3D nature of the alginate beads providing better spatial and mechanical support for matrix assembly. Hence the alginate system highlights the critical role of the 3D culture environment in promoting native-like conditions for the chondrocytes.

These cells display the three-dimensional round phenotype associated with chondrocytes *in vivo*. This suggests that formation and accumulation of a matrix through simulating 3D *in vivo* conditions is a crucial step that occurs in alginate encapsulation.

Similarly, cells isolated from alginate beads and re-embedded in agarose displayed a robust collagen VI and II staining in the outer matrix when cultured in ascorbate.

Metabolomic profiling found an upregulation in urea cycle and amino acid group metabolism pathways in compressed alginate-encapsulated chondrocytes. Distinct compression-induced changes in metabolomic profiles between the monolayer-agarose and alginate-released agarose-embedded chondrocytes indicated physiological changes in cell metabolism, likely to support differences in energy demands and matrix remodeling.

## Materials and Methods

### Chondrocyte Harvest and Culture

Primary human chondrocytes were harvested from n=6 Stage IV osteoarthritis patients (2 male and 4 female, age range 67-88 years old) under IRB approval using established methods [21]. Bovine chondrocytes were harvested from intact ankle joints of 18-22 month old cattle (n=6) obtained from a local abattoir. Chondrocytes were isolated by overnight digestion with Type I Collagenase (Gibco) at 2 mg/mL for 14 h at 37°C. Isolated chondrocytes were cultured in Dulbecco’s Modified Eagle’s medium (DMEM) supplemented with 10% Fetal Bovine Serum (FBS, Bio-Techne) and 10,000 I.U./mL penicillin and 10,000 μg/mL streptomycin (Sigma, hereby referred to as Complete media) in 5% CO2 at 37°C. Chondrocytes were passaged at 90% confluency (∼10 days) and used between passage 1-2. For monolayer studies, primary chondrocytes were seeded at a density of 160 cells/mm^2^ onto 25mm x 25mm Fisherbrand microscope coverslips and allowed to grow into a monolayer. The coverslips with the attached cells were placed in a 60mm x 15mm tissue culture dish containing Complete media and incubated for 0-7 days with 50μg/mL sodium L-ascorbate (Sigma) with the media changed every other day. Cells were analyzed at days 0, 3, and 7 for collagen production. Similarly, bovine chondrocytes were seeded at 160 cells/mm^2^ on microscope coverslips and cultured in Complete media with 50μg/mL sodium L-ascorbate for 7 days in 5% CO_2_ at 37°C and analyzed at days 0, 3, and 7.

Control cultures lacked sodium L-ascorbate in the media. Monolayers grown in Complete media lacking sodium L-ascorbate for 7 days at 37°C were used as a control.

### Alginate Encapsulation of Chondrocytes

Cells harvested from monolayers were encapsulated in alginate. For alginate encapsulation, cells were resuspended at a density of 4x10^6^ cells/mL in sterile filtered 1.2%, w/v, sodium alginate (Sigma). This solution was then slowly dispensed through a 22-gauge needle in a dropwise fashion into a 102mM CaCl_2_ solution. After near-instantaneous gelation, the alginate beads were allowed to crosslink for 15 min. The beads were then washed 4 times with sterile 150mM NaCl and once with complete media, before being cultured in 60 mm culture plates with complete media. The beads were maintained in complete media containing 50 *μ*g/mL sodium L-ascorbate for 0, 3 or 7 days with media changed every other day. Control cultures did not have ascorbate in the media. One group of alginate beads was cultured for 7 days with 50 *μ*g/mL sodium L-ascorbate for agarose encapsulation and mechanotransduction studies.

### Embedding in High-Stiffness Agarose

After 7 days in culture, the alginate beads were dissolved with a sterile EDTA-citrate buffer (150mM NaCl in 55mM sodium citrate with 50mM EDTA, pH6.8). The recovered cells were washed with PBS before mixing with low melting temperature agarose (Sigma Aldrich) at a final concentration of 4.5% w/w [22]. Agarose gels were placed in a 24-well tissue culture plate (Falcon) with 2 mL complete media containing 50 *μ*g/mL sodium L-ascorbate and cultured in 5% CO_2_ at 37°C for 3 or 7 days for metabolomic studies and histochemical analysis. The media was changed every other day. Monolayer cells embedded in agarose and cultured without ascorbate for 7 days served as controls.

### Cryosectioning, Immunocytochemistry, and Confocal Imaging

<u>Monolayer</u>: Primary human and bovine chondrocytes cells grown as monolayers on microscope coverslips were fixed in 4% Paraformaldehyde in 1X Phosphate Buffered Saline (PBS) for 10 min at room temperature followed by three 5 min washes of 1X PBS. Cells were permeabilized with 0.1% Triton and blocked with 10% normal goat serum in 1X PBS for 30 min at room temperature. Cells were then incubated with primary antibodies to both Collagen VI (Rabbit Polyclonal Anti-Collagen VI antibody, ab6588 from Abcam) and Collagen II (Mouse Monoclonal 2B1.5 to collagen II, ab186430, Abcam) in Triton X-100/1% Bovine Serum Albumin/1X PBS for 30 min at room temperature, followed by 0.1%Triton X-100/1% Bovine Serum Albumin/1X PBS for 1h at room temperature. After three 5 min washes with 1X PBS, cells were incubated with a mixture of secondary antibodies Donkey anti-Rabbit IgG H&L Alexa Fluor^R^488 (Abcam), Goat anti-Mouse IgG H&L Alexa Fluor^R^568 (Abcam) and Vibrant^TM^DyeCycle^TM^ Violet Stain for Nuclei (Invitrogen) in 0.1%Triton X-100/1% Bovine Serum Albumin/1X PBS for 1 h at room temperature. Cells were washed three times with 1X PBS and the coverslip was mounted on a glass slide with ProLong^TM^ Diamond Antifade Mountant (Invitrogen). Digital images were acquired on a Leica TCS SP8 confocal microscope and images were obtained with the Leica Application Suite Advanced Fluorescence software.

### Alginate-encapsulated chondrocytes

Alginate beads with encapsulated chondrocytes were spun down at 500 x g for 2 min, washed with 1X Phosphate Buffered Saline (PBS), and then fixed with 4% v/v Paraformaldehyde in 1X PBS for 15 min at room temperature, followed by a wash with 1X PBS. 100 μL of the alginate beads were cytospun onto single frosted adhesive slides (Tanner Scientific) using a Thermo Scientific Cytospin^TM^ 4Cytocentrifuge. Immunocytochemistry was then performed as described earlier.

### Agarose gels

Fixation and cryosectioning of agarose gels containing chondrocytes was performed using established methods with slight modification [23]. Agarose gels were fixed in 2 mL 4% v/v Paraformaldehyde – 1XPBS for 30 min at room temperature followed by 3 washes in 1X PBS. The gels were equilibrated overnight in 30% sucrose/1X PBS at room temperature followed by incubation in 2 mL of 50/50 30%sucrose/OCT solution for 2 h at room temperature. The gels were then cryopreserved in Tissue Tek OCT cryo-compound (VWR Scientific Products) at -80°C and 12 µm sections were mounted on charged slides (Tanner Scientific) for immunocytochemistry.

### Western Blot Analysis

Primary human and bovine chondrocytes were seeded at a density of ∼31000 cells/cm^2^ (2 mL media) in 6-well tissue culture plates (Falcon) and cultured in complete media containing 50μg/mL sodium L- ascorbate for 7 days in 5% CO_2_ at 37°C. Cells were collected at days 0, 3, and 7. Control cultures lacked ascorbate in the media.

Whole cell lysates were prepared from extracting proteins using a lysis buffer (0.1% Triton X-100/1XPBS containing a protease inhibitor). Protein concentration was measured with a Bicinchoninic acid (BCA) assay (Thermo Scientific Pierce^TM^ Rapid Gold BCA Assay Kit) and 8ug of total protein from each lysate was separated on a 4-15% SDS-polyacrylamide gel (Mini-PROTEAN® TGX™ Precast Protein Gels (BIO- RAD). After proteins were transferred to nitrocellulose membrane (BIO-RAD), membranes were blocked for 1 h in TBST containing 5% bovine serum albumin.

Blocked membranes were then incubated with primary antibody (1:2000 Rabbit Polyclonal Anti-Collagen VI antibody, Abcam) and incubated overnight at 4°C. Membranes were then washed four times with TBST for a total of 20 min, followed by incubation with Donkey anti-Rabbit IgG H&L Alexa Fluor^R^488 (1:2000 Abcam) for 1 h at room temperature. Loading controls were detected by running equal amounts of each lysate on a separate gel and probed with Rabbit Polyclonal Anti-Human, Anti-Bovine β actin (Rockland Immunochemicals). The nitrocellulose membranes were subsequently imaged using a Typhoon Trio Imager (GE).

### Chondrocyte Cyclical Compression and Metabolite Extraction

Agarose gels in PBS were placed in a custom-built bioreactor in 5% CO_2_ at 37°C, loaded initially to 5% strain, followed by cyclical compression with a 3.8% peak-to-peak strain amplitude at 1.1Hz for 30 min [22]. Immediately after compression, agarose gels were flash frozen for 1 min in liquid nitrogen. Frozen gels were crushed and placed in - 80°C before extraction with 1 mL of HPLC-grade methanol:acetone (70:30, Fisher Chemical). Pulverized gels with methanol:acetone were then vortexed every 5 min for 20 min and incubated overnight at -20°C for macromolecule precipitation. Samples were then centrifuged at 19000 x *g* for 10 min at 4°C (Sorvall Legend X1R). The supernatants were transferred to fresh tubes and vacuum dried in a Savant SC110 speed-vac. The resulting dried pellet was resuspended in 100 µl of 1:1 volume HPLC grade acetonitrile and water (Fisher Chemical) for further analysis by HPLC-MS.

### Untargeted Metabolomic Profiling and Statistical Analysis

Metabolites were detected via HPLC-MS (high performance liquid chromatography coupled to mass spectrometry) using an established protocol with a Cogent Diamond Hydride HILIC 120 x 2.1 mm column and Agilent 1290 UPLC system [24]. Raw data were acquired in positive ion mode. Initially, we used MS-Convert to re-format raw files, and XCMS Online for feature detection, retention time correction, and alignment [25].

Metaboanalyst was used to the normalize the data and to run the following statistical analyses: principal component analysis (PCA), partial least squares-discriminant analysis (PLS-DA), ANOVA (for multiple group comparisons), t-test (for two group comparisons), hierarchical cluster analysis (HCA), heatmap and volcano plot analysis (for two group comparisons). Significant metabolomic features resulting from volcano plots and heatmap clusters were used to run the pathway analysis on MetaboAnalyst’s “Functional Analysis” feature.

## Results

### Collagen VI and II expression is up regulated after culture in ascorbate and increased through day 7

To study the expression of types VI and II collagen in monolayers and alginate beads we cultured primary human chondrocytes in 50μg/mL sodium L- ascorbate for 3 or 7 days in 5% CO_2_ at 37°C with no ascorbate media serving as a baseline control (day 0, Fig. 1). Collagen synthesis relies on the hydroxylation of proline and lysine, which are key components of the collagen triple helix structure. During hydroxylation, ascorbic acid acts as a critical cofactor for the enzymes, prolyl hydroxylase and lysyl hydroxylase which catalyze the addition of hydroxyl groups onto proline and lysine to convert them into hydroxyproline and hydroxylysine respectively [26]. To evaluate the localization of collagen VI and II within different culture environments, we employed immunofluorescence across three different conditions: chondrocytes grown in monolayers, alginate-encapsulated chondrocytes, and chondrocytes that had been released from alginate and subsequently embedded in agarose.

**Figure 1.**
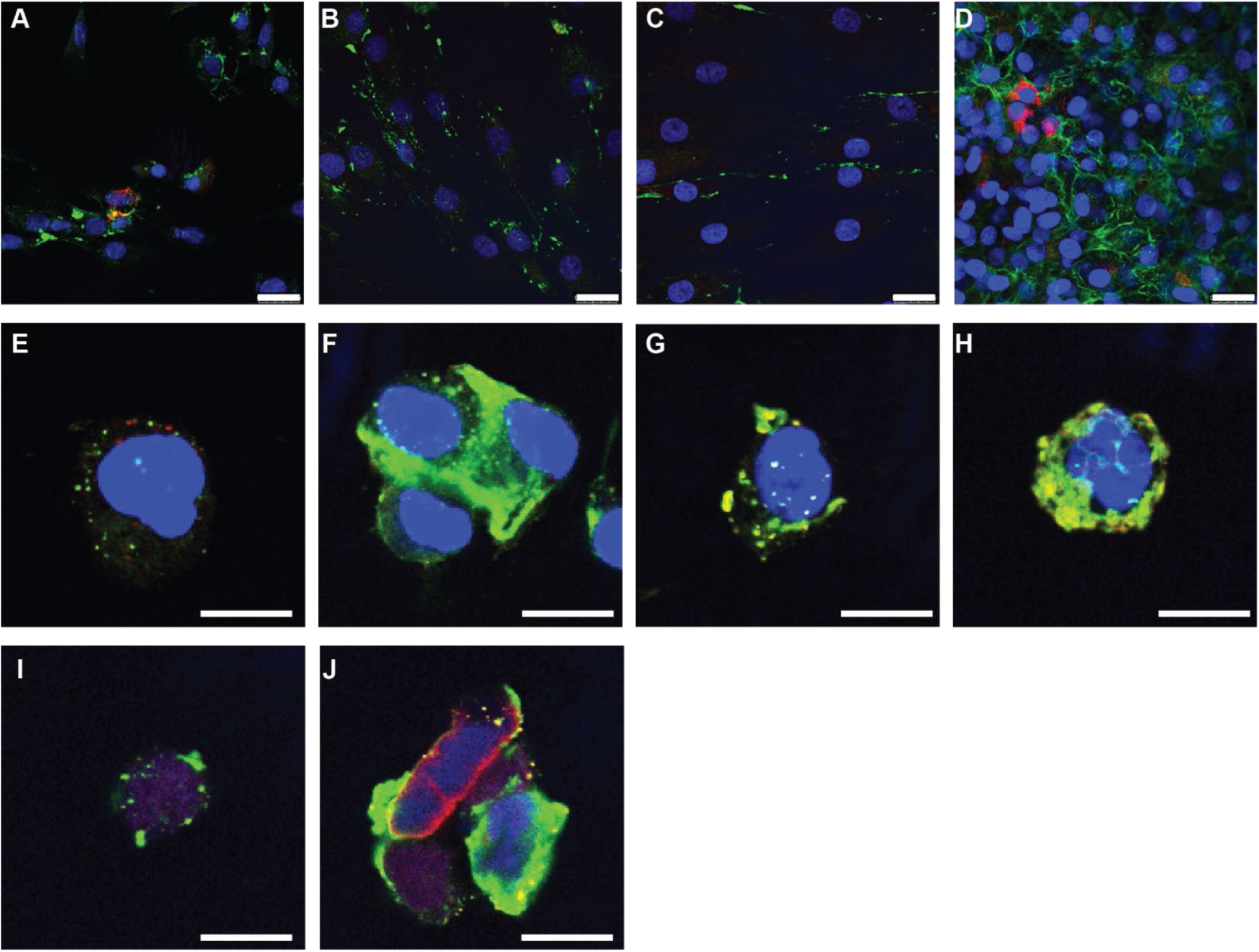
Collagen expression in primary human chondrocytes. Primary human chondrocytes were cultured and stained with antibodies to Collagen VI (green), Collagen II (red) and Nuclear Marker Vibrant^TM^DyeCycle^TM^ Violet Stain before imaging at 63X. Monolayer-expanded human primary chondrocytes showed a progressive increase in collagens VI and II when cultured (**A**) at day 0, (**B**) day 3 with ascorbate (**C**) day 7 no ascorbate control and (**D**) day 7 with ascorbate. (**E**) Alginate-encapsulated chondrocytes at day 0, (**F**) at day 3 with ascorbate, (**G**) day 7 without ascorbate and (**H**) day 7 with ascorbate in the medium. Sections of chondrocytes released from alginate and re-embedded in agarose stained for Collagens VI and II expression in the pericellular matrix (**I**) when cultured without ascorbate and (**J**) with ascorbate for 7 days. Scale bar represents 25 µm in panels A-D and 12.5 µm in E-J.

Using double immunofluorescence staining for collagens VI and II, we observed that compared to day 0 (Fig. 1A), primary human chondrocytes when cultured as a monolayer in ascorbate showed a progressive increase in collagen production by day 3 (Fig. 1B). This increase in collagen VI and II production was most noticeable at day 7 (Fig. 1D). In contrast, monolayer controls cultured without ascorbate displayed considerably less collagen production at day 7, emphasizing the role of ascorbate in collagen synthesis (Fig. 1C). Given these findings, we hypothesized that culturing primary human and bovine chondrocytes in alginate would mimic *in vivo* physiological conditions and, in the presence of ascorbate would help synthesize a cell-directed matrix formation. Hence, we encapsulated primary human chondrocytes in 1.2% sodium alginate and cultured them in ascorbate for varying lengths of time.

Similar to the trends observed in monolayer cultures, alginate encapsulated chondrocytes demonstrated time-dependent increase in collagen synthesis. Compared to the baseline day 0 (Fig. 1E), alginate-encapsulated chondrocytes synthesized higher amounts of collagen VI and to a lesser extent collagen II at day 3 (Fig. 1F) and day 7 (Fig. 1H). As expected, when cultured without ascorbate, alginate-encapsulated chondrocytes produced less collagen (Fig. 1G). Interestingly, we observed that chondrocytes cultured with ascorbate within the alginate matrix produced more collagen VI which is found in the pericellular matrix than collagen II. We noticed that the chondrocytes cultured in alginate beads had clearly formed chondrons (*i.e.* chondrocytes with a shell-like PCM) that included more collagen VI near the perimeter of the cell in a rounded 3D morphology. Collagen VI is a key component of the PCM playing a pivotal role in structural integrity, cell signaling, and cell-matrix interactions.

The high levels of Collagen VI synthesized under these conditions, and its localization around the chondrocytes specifically in the PCM suggests the formation of distinct chondrons. This observation supports our hypothesis that culturing chondrocytes in alginate and ascorbate would mimic *in vivo* conditions. This was in stark contrast to the more diffuse pattern of secreted collagen VI produced by monolayer chondrocytes which lacked the spherical PCM.

To further explore if these chondrons formed within the alginate culture system would retain their collagen expression and 3D phenotype upon released from alginate, chondrons were released from alginate beads and re-embedded in 4.5% physiologically stiff agarose and cultured for 7 days supplemented with ascorbate. Alginate released, agarose-embedded chondrocytes sustained collagen production in the presence of ascorbate when compared to the no ascorbate control (Fig. 1I-J) and retained their 3D round phenotype. This finding reiterates that culturing chondrocytes with ascorbate in an alginate system closely mimics *in vivo* conditions, fostering not only collagen production but also maintaining their *in vivo* structural characteristics.

We further wanted to evaluate collagen expression in healthy chondrocytes cultured with ascorbate and used a bovine model. When grown as monolayers, compared to the day 0 baseline control (Fig. 2A) bovine chondrocytes showed a progressive increase in collagen VI and II production from day 3 (Fig. 2B) to day 7 (Fig. 2D). When cultured without ascorbate at day 7 (Fig. 2C), bovine chondrocytes still produced collagen albeit to a lesser extent than in the presence of ascorbate. We then encapsulated the bovine chondrocytes in alginate to mimic a 3D environment and cultured them in ascorbate. As expected for healthy chondrocytes, these alginate-encapsulated bovine chondrocytes expressed collagen VI and II at day 0 (Fig. 2E), which substantially increased by day 3 (Fig. 2F) and day 7 (Fig. 2H) compared to the 7 day no ascorbate control (Fig. 2G). Notably, when cultured in ascorbate, bovine chondrocytes in alginate beads proceeded to form a shell-like pericellular matrix that surrounds the cell, similar to the primary human chondrocytes in alginate cultures.

**Figure 2.**
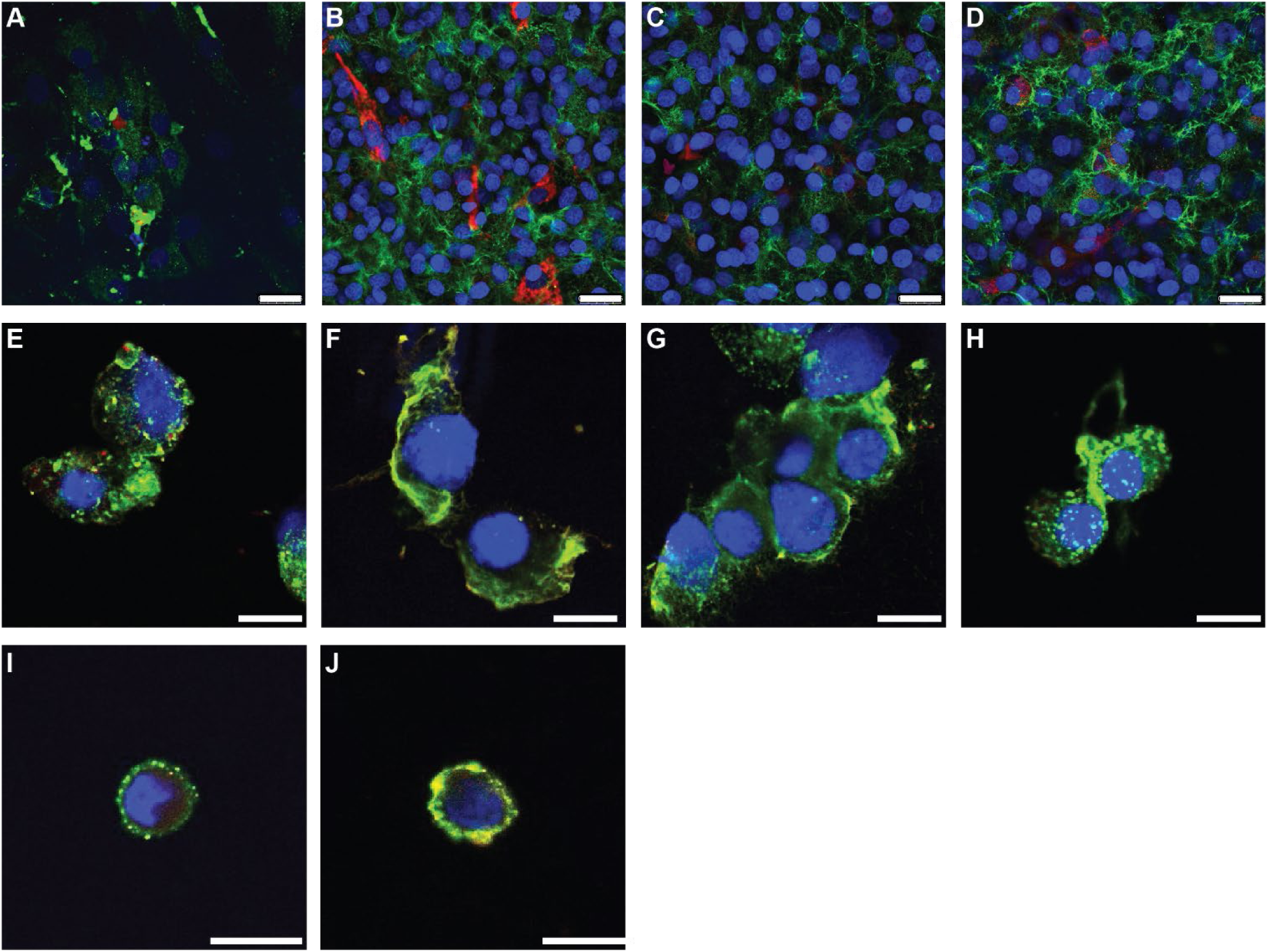
Expression of Collagens VI and II in bovine chondrocytes. Representative 63x images of bovine chondrocytes labeled with antibodies to Collagen VI (green), Collagen II (red) and Nuclear Marker Vibrant^TM^DyeCycle^TM^ Violet Stain. Panels (**A-D**) are monolayer expanded bovine chondrocytes from (**A**) day 0, (**B**) day 3 cultured with ascorbate, (**C**) day 7 no ascorbate control and (**D**) day 7 cultured with ascorbate. Panels (**E-H**) represent alginate beads cultured (**E**) at day 0, (**F**) day 3 with ascorbate, (**G**) day 7 no ascorbate culture and (**H**) day 7 ascorbate. Immunofluorescence staining of agarose sections depicting chondrocytes released from alginate beads and re-embedded in agarose and (**I**) cultured without ascorbate and (**J**) with ascorbate for 7 days. Scale bar represents 25 µm.

To examine the stability of collagen production in agarose, we released the alginate encapsulated bovine chondrocytes and re-embedded them in physiologically stiff agarose and cultured them with ascorbate. After 7 days, alginate-released, agarose-embedded bovine chondrocytes displayed robust collagen expression in the pericellular matrix when cultured with ascorbate (Fig. 2I) and to a lesser extent in the absence of ascorbate (Fig. 2J). These results confirm that culturing bovine chondrocytes in alginate results in the formation of a well-defined matrix synthesis.

### Collagen VI protein expression is increased in monolayer cultured in ascorbate for 7 days

To assess collagen VI protein levels, we conducted western blot analysis from extracts of both primary human and bovine chondrocytes cultured as monolayers in the presence and absence of ascorbate over a 7-day period. Chondrocytes allowed to adhere and grow for 24 h without ascorbate served as day 0 baseline controls. We used a rabbit anti-ColVI polyclonal antibody for detection. We noticed that collagen VI protein levels increased in cultured primary human chondrocytes from day 0 (Fig. 3A, lane 1) to day 7 without ascorbate (Fig. 3A, lane 2) and when supplemented with ascorbate (Fig. 3A, lane 3). Similarly, bovine chondrocytes cultured at day 0 (Fig. 3B, lane 4) had less collagen compared to chondrocytes cultured for 7 days without ascorbate, but substantial increases at day 7 cultured with ascorbate (Fig. 3B, lanes 5-6).

**Figure 3.**
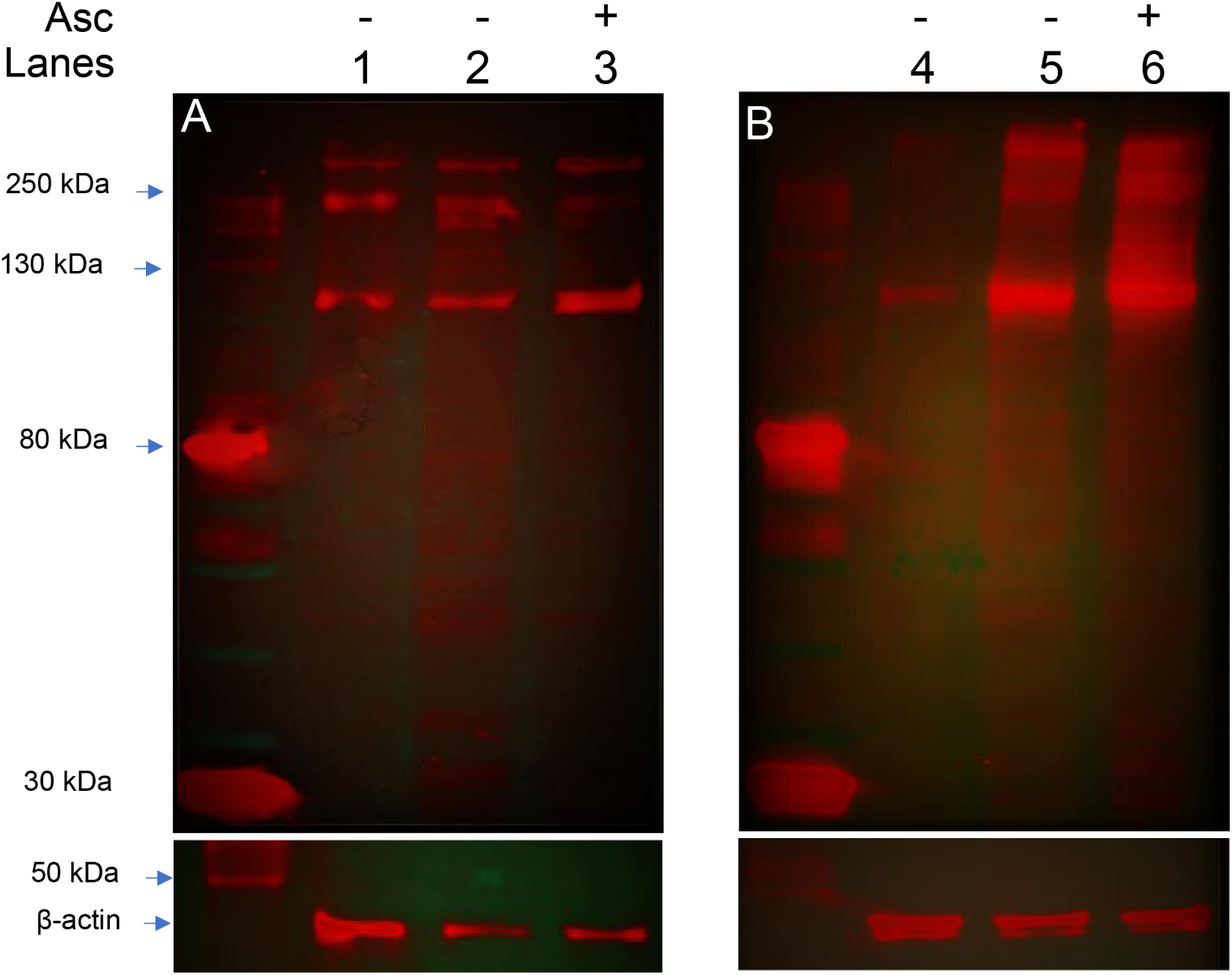
Ascorbate increases Collagen VI expression. Western blot analysis of primary human and bovine chondrocytes. Monolayer cell lysates from **(A)** primary human chondrocytes and **(B)** bovine chondrocytes were probed with antibody to Collagen VI. Lanes 1 and 4 represent cell lysates from day 0, lanes 2 and 5 represent cell lysates from monolayers cultured for 7 days without ascorbate (Asc). Cell lysates from monolayers cultured with ascorbate (Asc) for 7 days are shown in lanes 3 and 6. β-actin was used as loading control.

When analyzing whole cell lysates under reducing conditions, two major bands were detected, one at ∼ 250 -300 kDa and another ∼ 120 kDa. These bands represent the different components of the collagen VI triple helix: the larger band corresponds to the bigger α3(VI) chain and the smaller band most likely corresponds to partially processed α1(VI) and α2(VI) chains. These low molecular weight bands could also be small degradation or post-translationally modified products. We did see a smeared appearance of the bands in the bovine chondrocytes, possibly due to the abundance of collagen VI produced in these healthy chondrocytes and incomplete denaturation under our reducing conditions.

### Distinct metabolomic profiles are observed in primary human and bovine chondrocytes when subjected to cyclical compression

To investigate whether alginate pre-encapsulation influences cellular mechanotransduction, alginate-encapsulated chondrocytes were embedded in physiologically-stiff agarose and subjected to mechanical loading before metabolite extraction and metabolomic profiling. Monolayer cultured chondrocytes embedded in agarose were treated similarly as controls.

Untargeted metabolomic profiling detected 2678 metabolite features in primary human chondrocytes and 3322 in bovine chondrocytes (Supplemental Tables 1-2).

Metabolite features identified as significant by ANOVA (FDR-adjusted p-values<0.05) were used for further analyses. To assess overall variation between the different experimental groups, we used unsupervised Principal Components Analysis (PCA) and Hierarchical Clustering Analysis (HCA). PCA found moderate separation with more than 50% of the total variance associated with the first 3 components comparing alginate-encapsulated agarose-embedded and monolayer cultured agarose-embedded primary human chondrocytes (Fig. 4A). This separation may suggest that pre-encapsulation in alginate may influence metabolic pathways potentially affecting cellular mechanotransductive response. There was moderate separation by HCA between alginate-encapsulated and monolayer-cultured agarose-embedded primary human chondrocytes under uncompressed conditions (Fig. 4B). Compared to the primary human chondrocytes, PCA found slightly less separation between alginate-encapsulated and monolayer cultured agarose-embedded uncompressed bovine chondrocytes (Fig. 4C). Compared to primary human chondrocytes, bovine chondrocytes had less distinct clustering of HCA identified metabolites suggesting that they exhibit less metabolic variability reflecting species-specific metabolism (Fig. 4D).

**Figure 4.**
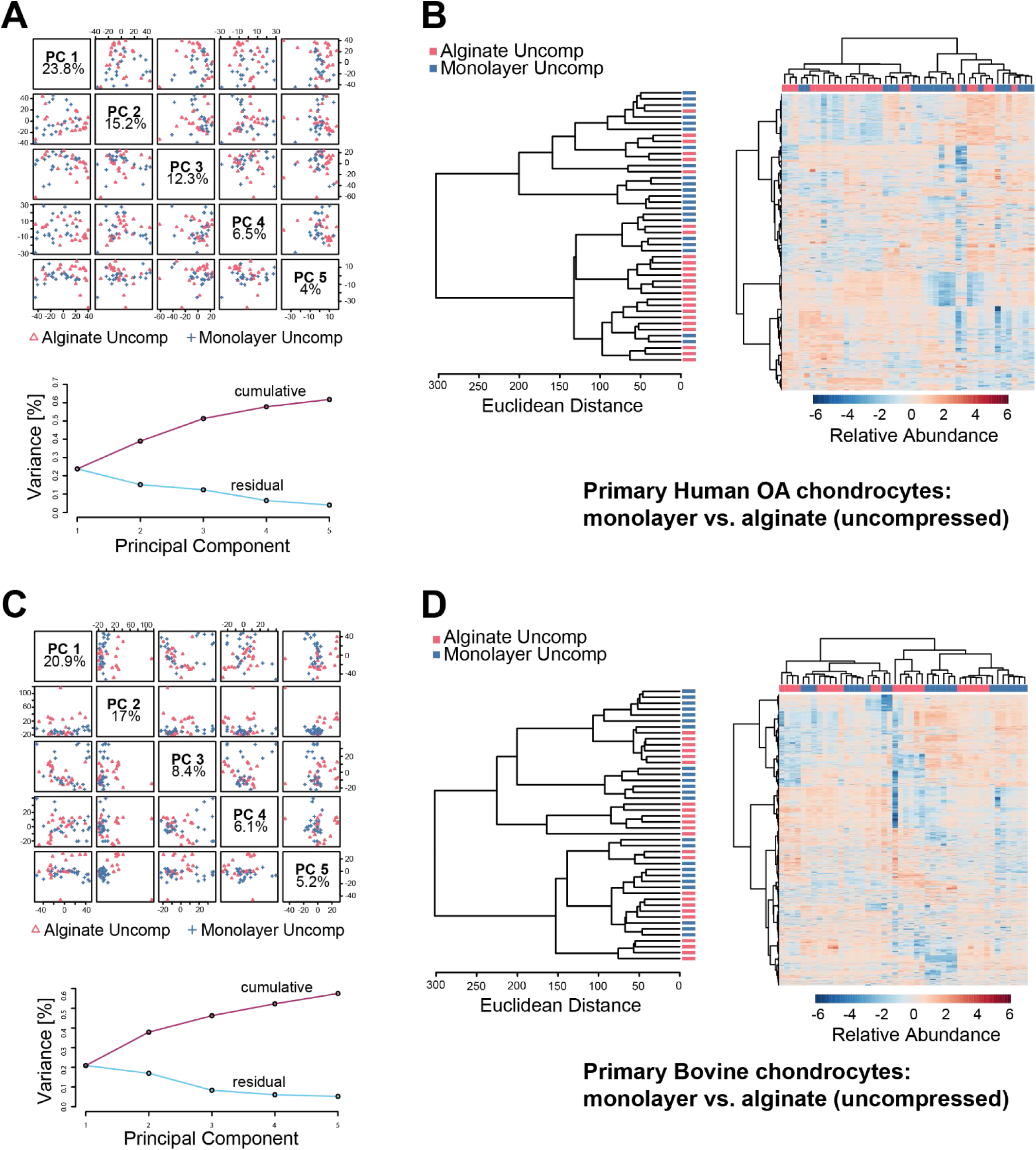
Alginate pre-culture drives differences in baseline metabolomic profiles in uncompressed chondrocytes. (A) For primary human OA chondrocytes principal components analysis finds substantial differences between monolayer controls and alginate pre-cultured samples with more than 50% of the overall variance associated with the first 3 components. (B) Clustering analysis finds good discrimination between monolayer and alginate pre-cultured samples of primary human chondrocytes. (C) For bovine chondrocytes, principal components analysis finds differences between monolayer and alginate pre-culture samples. (D) Hierarchical clustering finds minimal discrimination between monolayer and alginate pre-cultured bovine chondrocytes.

We then evaluated the effects of cyclical compression on both alginate-released- agarose-embedded and monolayer-agarose-embedded chondrocyte metabolism. PCA captured 50% of the variance within the first 3 components indicating variation in metabolomic profiles due to mechanical loading (Fig. 5A). Cluster analysis of the top 25 metabolites did not find distinct clustering between the alginate and monolayer cultured chondrocytes, suggesting donor dependent heterogeneity and differential regulation of pathways and metabolites in response to compression (Fig. 5B). Conversely, for bovine chondrocytes subjected to same compression conditions, 44% of variation was seen in PCA reflecting different distribution of metabolic changes (Fig. 5C) and HCA showed perfect clustering of metabolites with respect to alginate and monolayer cultured chondrocytes in response to compression (Fig. 5D). This clear separation indicates that unlike human chondrocytes, bovine chondrocytes exhibit a more uniform response to cyclical compression with each culture condition (alginate vs monolayer).

**Figure 5.**
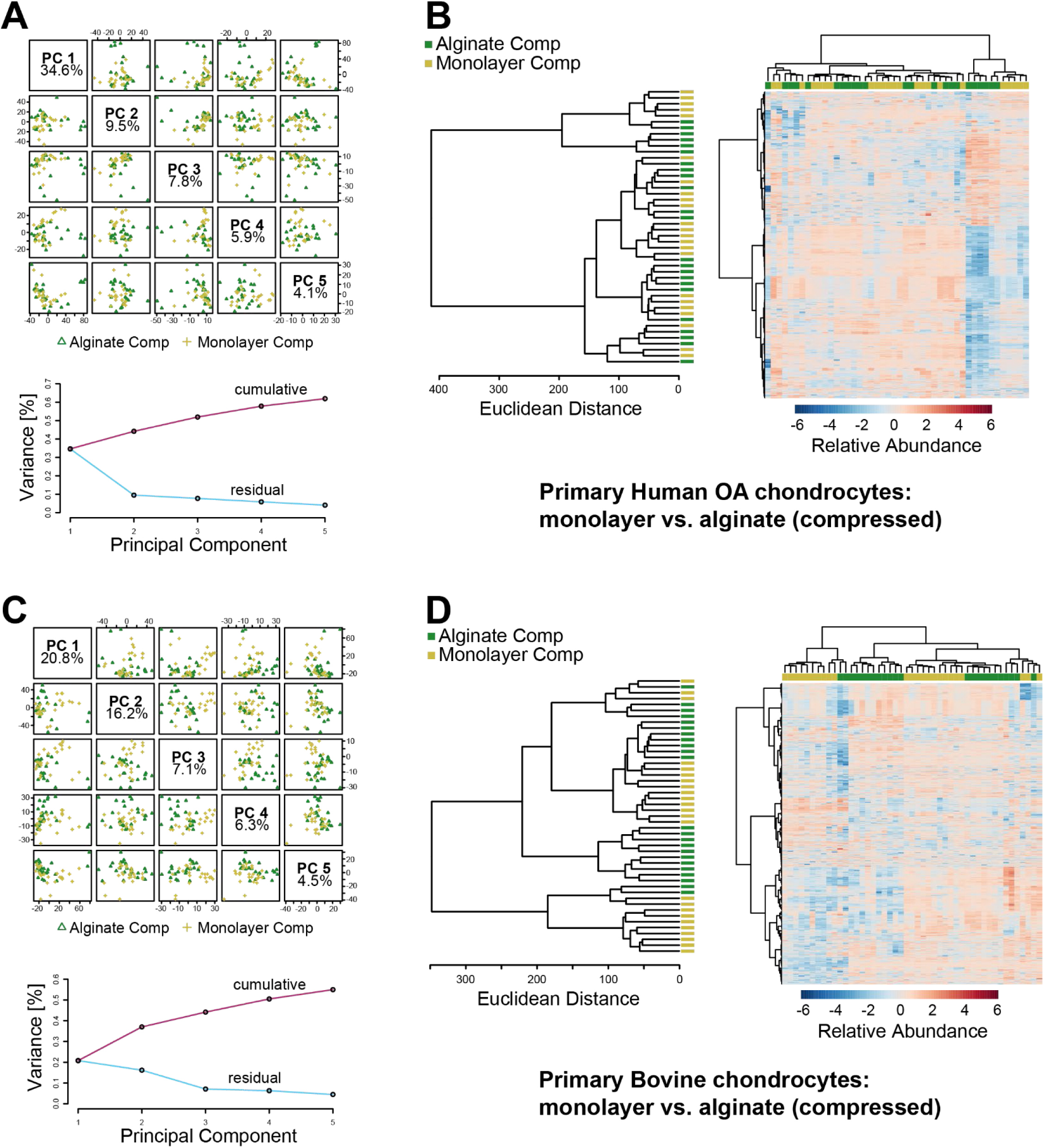
Alginate-encapsulated chondrocytes exhibit differences in compression-induced metabolomic profiles compared to monolayer-encapsulated chondrocytes. Metabolomic profiles from primary human chondrocytes shown in panels (A) and (B). Metabolomic profiles from bovine chondrocytes shown in panels (C) and (D).

We further compared the effects of compression on alginate-released-agarose-embedded primary chondrocytes and monolayer-agarose-embedded primary chondrocytes (Fig.6). PCA and clustering displayed minimal overlap between the compressed and uncompressed alginate-released-agarose-embedded chondrocytes. The variation in the data set for the first three components was higher than 50%, demonstrating the importance of compression for driving changes in the chondrocyte metabolome. Similarly, heat map analysis revealed compression-induced differential changes in the metabolites obtained from both the groups. Hierarchical clustering also revealed that some of the clusters of metabolites were upregulated in the compressed alginate groups (light green) and comparatively downregulated in the uncompressed alginate group (light pink) indicating the heterogeneity in the response of primary human chondrocytes to compression (Fig. 6A). This response can be attributed to sexual dimorphism of the donor cells as well as to the physiological condition of the harvested chondrocytes. Upon examining the effect of compression on monolayer-agarose-embedded chondrocytes we found moderate separation (44.5%) between the first three components. HCA and heat map analysis further revealed differences in regulation of metabolites between the compressed and uncompressed monolayer cultured chondrocytes, validating their response to compression (Fig. 6B).

**Figure 6.**
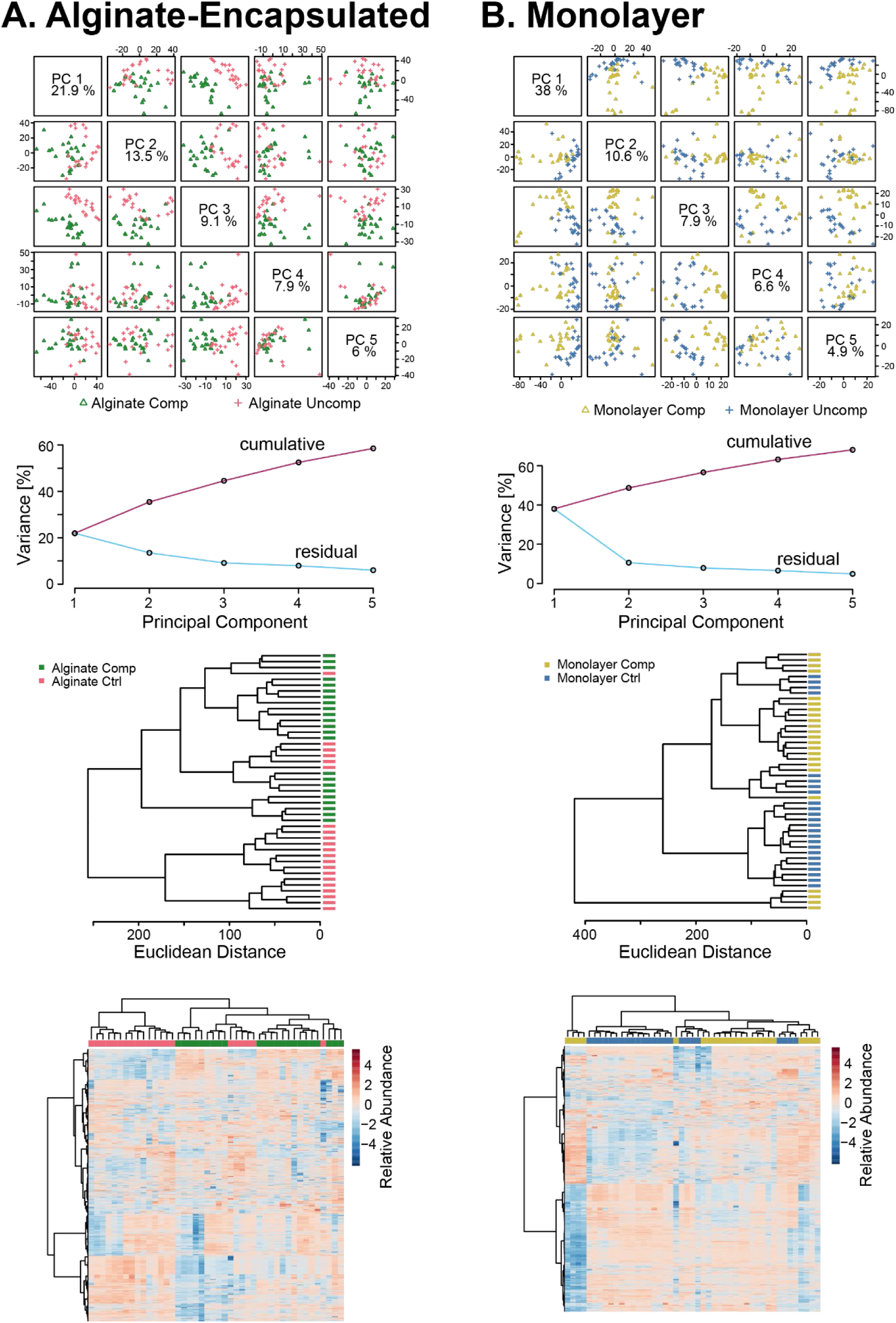
Comparison of compression-induced metabolomic profiles between alginate-encapsulated (A) and monolayer (B) primary human chondrocytes. Principal components analysis (top) finds that both systems exhibit differences between compression-induced changes in metabolomic profiles. Clustering and heatmaps show differences in compression-induced metabolomic profiles (middle and bottom).

Unsupervised clustering showed limited variability between compressed and uncompressed bovine chondrocytes cultured in alginate. HCA and heat map analysis did not find distinct clustering of the metabolites in the alginate compressed and uncompressed group (Fig. 7A). Similarly, not much difference was observed in the compressed and uncompressed chondrocytes grown as monolayers and embedded in agarose. Distinct metabolomic profiles were not observed by HCA and heat map analysis (Fig. 7B). These findings imply that for bovine chondrocytes the transition from uncompressed to compressed, irrespective of the culture system, may not show distinct changes in the metabolic profiles that can be detected by untargeted analysis.

**Figure 7.**
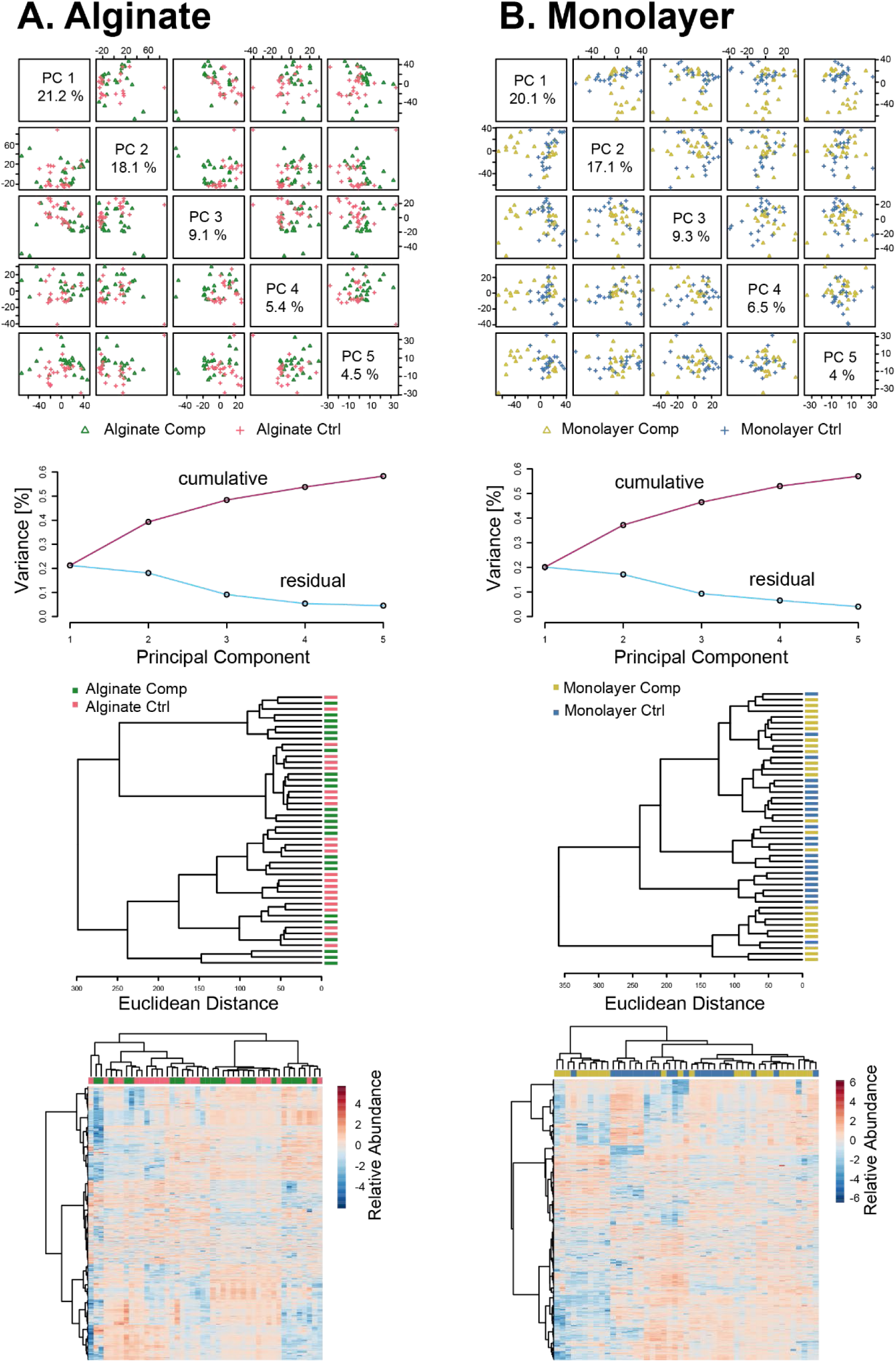
Comparison of compression-induced metabolomic profiles between alginate-encapsulated (A) and monolayer (B) bovine chondrocytes. Principal components analysis (top) finds that both systems exhibit differences between compression-induced changes in metabolomic profiles. Clustering and heatmaps show differences in compression-induced metabolomic profiles (middle and bottom).

In primary human chondrocytes, hierarchical clustering of the top 50 metabolites displayed distinct clusters of metabolites that were upregulated in the compressed alginate cultured-agarose-embedded cells (Fig. 8A, green) and downregulated in the uncompressed (Fig. 8A, pink). Interestingly, in the monolayer cultured-agarose-embedded chondrocytes hierarchical clustering of the top 50 metabolites revealed downregulation of metabolites in the compressed (Fig. 8B, yellow) and an upregulation in the uncompressed groups (Fig. 8B, blue). This suggests that compression in an alginate-agarose environment activates specific metabolic pathways reflecting the cells’ response to stress. These findings highlight the impact of initial culture environment on chondrocyte metabolism and mechanotransduction pathways.

**Figure 8.**
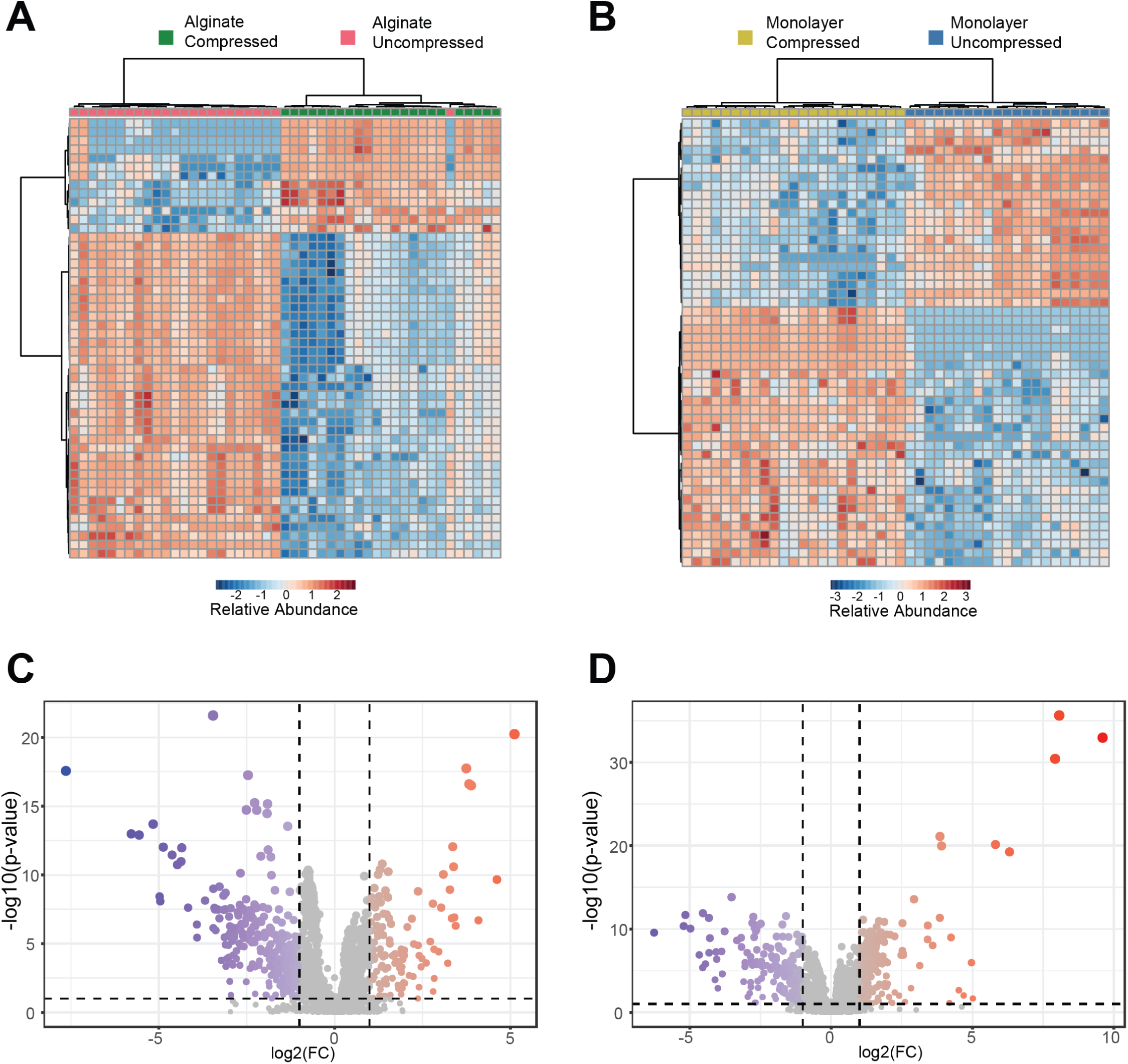
Comparison of mechanotransduction between alginate and monolayer systems. (A) Top 50 metabolites different between compressed and uncompressed controls in alginate system. (B) Top 50 metabolites different between compressed and uncompressed controls in monolayer system. (C) Volcano plot of compression-induced metabolites in alginate system. (D) Volcano plot of compression-induced metabolites in monolayer-agarose system.

We further utilized volcano plot analysis (Fig. 8C) to find metabolite features that differ between the alginate cultured and monolayer cultured chondrocytes encapsulated in agarose gels and subjected to cyclical compression. 120 metabolites were significantly upregulated in the alginate cultured-agarose-embedded chondrocytes subjected to compression and 354 metabolites were downregulated. Pathway analysis of upregulated metabolites indicated an enrichment of pathways related to branched- chain amino acid degradation, urea cycle/amino group metabolism, steroid hormone biosynthesis, lysine metabolism and hormone metabolism. Downregulated pathways mapped to aminosugars and pyrimidine metabolism pathways in the alginate cultured cells.

For monolayer-cultured chondrocytes embedded in agarose and subjected to compression, we identified 192 significantly upregulated metabolites and 185 downregulated metabolites (Fig. 8D). Upregulated pathways included fatty acid metabolism, saturated fatty acid oxidation, and tryptophan metabolism pathways. This shift toward fatty acid and tryptophan metabolic pathways could suggest a cellular strategy to meet increased energy demands or modulate stress responses under compression in monolayer cultures. Downregulated metabolites represented pathways such as N-glycan biosynthesis, butanoate metabolism, and urea cycle/amino group metabolism. Interestingly, some of the metabolites that were found in high abundance in the alginate cultured-agarose-embedded compressed group, such as hippuric acid, were at lower concentration in the monolayer-cultured agarose-embedded compressed group. Hippuric acid has been noted in studies as a potential biomarker, where a study by Rushing *et al* showed that levels of hippuric acid were increased in urine samples of OA progressors, suggesting the importance of hippuric acid as a metabolite in distinguishing OA progression [27]. The observed difference in hippuric acid levels between culture conditions in our study could imply that alginate-cultured chondrocytes more closely mimic physiological conditions relevant to OA, providing a potentially valuable model for studying disease progression and metabolic adaptations to mechanical stress.

### Distinct compression-induced metabolic pathways are associated with monolayer and alginate-cultured chondrocytes and vary between primary human and bovine chondrocytes

Median heat map analysis of co regulated metabolites identified 9 different clusters in primary human chondrocytes and 8 clusters in bovine chondrocytes. We detected many pathways that were upregulated in the uncompressed alginate cultured- agarose-embedded group including amino acid, urea cycle, Vitamin B6 and B3 metabolism, glycerophospholipid metabolism, and energy cycle metabolism (Fig. 9A, Clusters # 3 and 4, dark blue).

**Figure 9.**
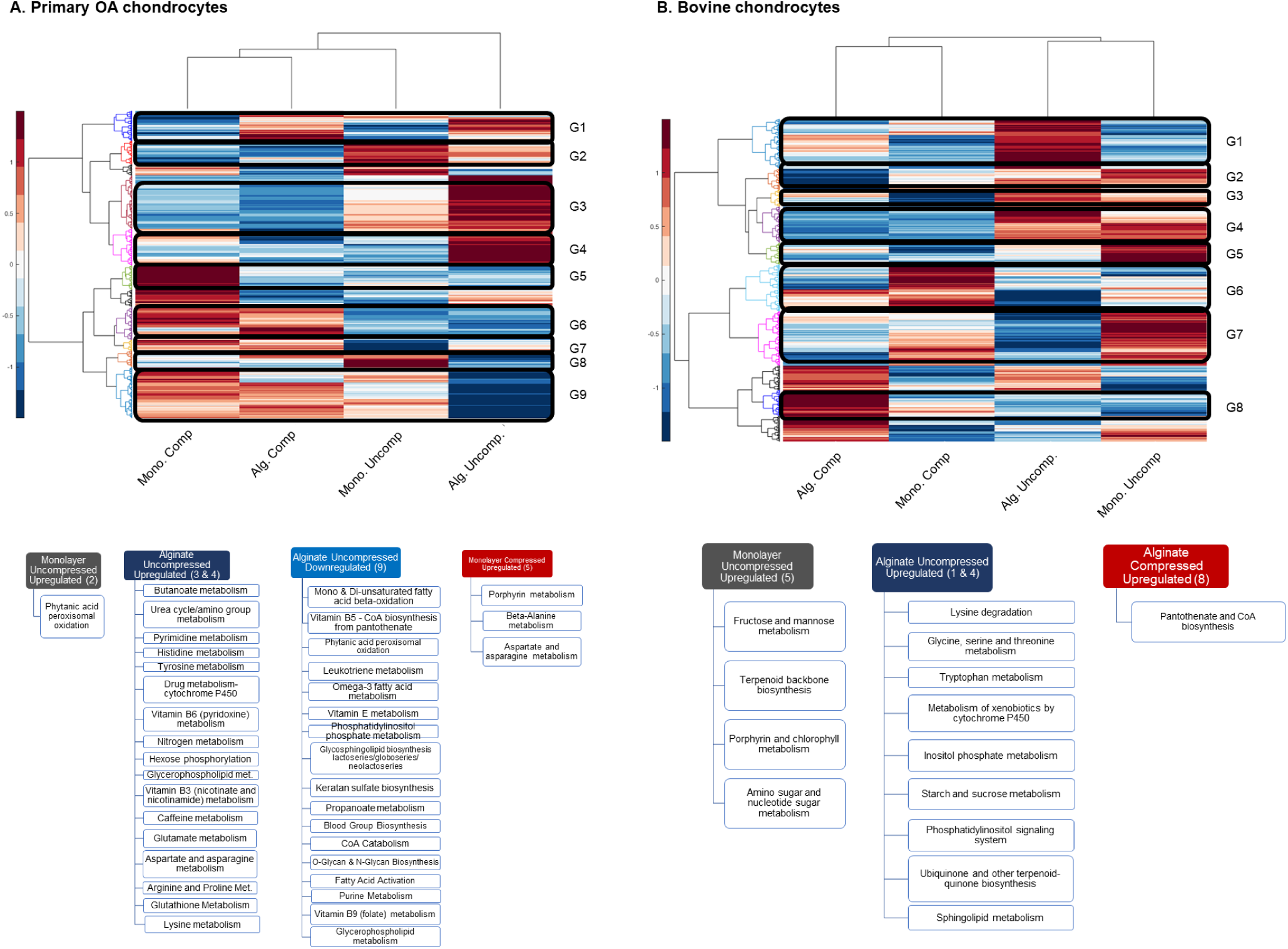
Summary of culture-related and compression-induced pathways for (A) primary OA chondrocytes and (B) bovine chondrocytes. Pathways determined from clusters of metabolites shown in top heatmaps.

Glycine and proline are major components of collagen, and we detected upregulation of several amino acids including arginine, proline, lysine, glutamate, histidine, tyrosine, aspartate and asparagine metabolism. Proline can be synthesized *de novo* from glutamate through the intermediate pyrroline-5-carboxylate (P5C), which is then converted to proline by P5C reductase [28]. The finding of upregulation of this pathway suggests that uncompressed alginate-cultured chondrocytes actively support collagen synthesis and matrix integrity.

Vitamin B6 is a critical cofactor involved in diverse biochemical reactions that regulate cellular metabolism including amino acid, glucose, and lipid metabolism, which are vital for cellular energy and overall metabolic regulation [29], whereas vitamin B3 (also known as niacin) plays a central role in energy metabolism [30].

In contrast, beta-oxidation of fatty acids in peroxisomes was uniquely upregulated in the uncompressed monolayer-cultured, agarose-embedded chondrocytes (Fig. 9A, Cluster # 2, dark green). Beta-oxidation within peroxisomes facilitates the breakdown of very long-chain fatty acids, which are converted into shorter-chain fatty acids that can enter mitochondrial beta-oxidation for energy production. This upregulation of fatty acid beta-oxidation in monolayer-cultured chondrocytes may reflect a reliance on lipid metabolism and alternative energy pathways [31].

Pathways related to porphyrin and amino acid metabolism, as well as beta alanine, aspartate and asparagine metabolism were upregulated in compressed monolayer group (Fig. 9A, Cluster # 5, red). The upregulation of amino acid metabolism pathways suggests that compressed monolayer chondrocytes may rely on these pathways to provide substrates for protein synthesis, energy production, and cellular repair processes.

Downregulated metabolic pathways for the uncompressed alginate group were mainly associated with fatty acid oxidation, keratan sulfate biosynthesis, glycan biosynthesis, glycerophospholipid metabolism and purine metabolism (Fig. 9A, Cluster # 9, blue). This decrease in fatty acid oxidation may suggest a reduced energy need from lipid sources, possibly due to the differences in the energy consumption in loading versus unloading states. The downregulation of keratan sulfate and glycan biosynthesis pathways points to a decreased focus on matrix remodeling in the uncompressed state. Similarly, the downregulation of glycerophospholipid metabolism suggests limited or basal membrane remodeling indicative of unloaded cells. Lastly, the downregulation of purine metabolism in the uncompressed alginate group may reflect a more quiescent metabolic profile in cells not subjected to the energetic and structural demands of compression. These distinct shifts between compressed and uncompressed conditions highlight the adaptive responses of chondrocytes in relation to energy utilization and structural components synthesis.

Upregulated pathways in the uncompressed alginate cultured-agarose- embedded bovine chondrocytes included lysine degradation, metabolism of amino acids glycine, threonine, serine and tryptophan, inositol phosphate metabolism, sphingolipid metabolism, and the phosphatidyl inositol signaling system, showcasing an active metabolic profile in the absence of loading (Fig. 9B, Cluster # 1, 4, blue). Inositol phosphates play key roles in cellular signaling pathways, including cell proliferation, differentiation, and stress responses and the phosphatidyl inositol signaling system is integral to various signaling cascades, including those involved in membrane dynamics and cellular responses to environmental changes. The energy related pathways such as starch and sucrose metabolism and ubiquinone biosynthesis were also upregulated in this cluster.

Pathways that were upregulated in the bovine uncompressed monolayer-cultured group included sugar metabolism for energy production, terpenoid backbone synthesis and electron transport and amino sugar metabolism (Fig. 9B, Cluster #5, dark green).

The terpenoid pathway is responsible for synthesizing isoprenoid units that form the backbone of many essential biomolecules, including cholesterol, steroids, and is essential for synthesizing ubiquinone (coenzyme Q), which plays a critical role in the mitochondrial electron transport chain. Furthermore, amino sugar metabolism was upregulated, which plays a vital role in producing amino sugars used for building glycosaminoglycans (GAGs). The upregulated pathway in the alginate compressed cluster was related to an energy pathway, Panthotenate and CoA biosynthesis (Fig. 9B, Cluster # 8, red). Panthotenate is a key precursor to Coenzyme A biosynthesis, which is an essential cofactor and plays an important role in synthesis of phospholipids, fatty acids and the TCA cycle [32].

## Discussion

The local microenvironment governs and is fundamental in regulating the structure-function relationships of chondrocytes within the articular cartilage. Mechanical signals play important roles in maintaining homeostasis of the articular cartilage extracellular matrix, and dysregulation can impair normal physiologic processes leading to chondrocyte dysfunction and OA progression [33]. The objective of this work was to develop a physiologically relevant culture system for chondrocytes to produce a robust pericellular matrix for studying chondrocyte mechanotransduction.

Ascorbic acid is an essential cofactor for collagen formation and its requirement is mainly attributed to its role in prolyl hydroxylation. Hydroxyproline is important for the helical structure formation of collagen fibers and hydroxylysine plays a critical role in collagen crosslinking. Hence, we used ascorbate in our culture medium to ensure production of collagen. Previous studies [20] show that ascorbic acid in culture medium enhances collagen production, and we also found this in the present study for both primary human and bovine chondrocytes. We observed that both primary human and bovine chondrocytes in monolayer culture supplemented with ascorbate showed a pronounced diffuse pattern of secreted collagens VI and II. This diffuse pattern reflects the less structured and planar environment of monolayer culture, which lacks the physical constraints and spatial cues of a 3D matrix. In contrast, both primary human and bovine chondrocytes cultured in alginate developed a robust three-dimensional round phenotype similar to *in vivo* chondrons and produced more collagen VI compared to monolayer. To further corroborate our immunofluorescence findings, we performed western blot analysis, which confirmed a substantial increase in collagen VI production in both human and bovine chondrocytes cultured with ascorbate.

In healthy joints, primary articular chondrocytes maintain physiological and metabolic homeostasis which is altered in osteoarthritic chondrocytes. This pathological shift includes cartilage remodeling characterized by degradation of the articular cartilage, disruption of the architecture, and a progressive deterioration of biomechanical properties. Changes within the chondrocytes include but are not limited to disturbed mitochondrial function, changes in energy requirements, altered cell signaling, and chondrosenescence [34].

We are beginning to understand some of the biochemical events that are involved in mechanotransduction and subsequent modulation of the chondrocyte phenotype. Cell-matrix interactions are crucial to mechanical sensing and this study shows that alginate-agarose constructs provide an improved model for studying chondrocyte mechanotransduction. Here, we identify conditions for chondrocytes to form an endogenous pericellular matrix and assess biochemical pathways affected by cyclical compression.

Metabolomic profiling of primary human chondrocytes precultured in alginate showed differentially regulated pathways related to energy metabolism, lipid metabolism, and amino acid metabolism compared to chondrocytes cultured as monolayer and embedded in agarose when subjected to cyclical compression. Volcano plot analysis found upregulated metabolite features mapped to branched chain amino acid (BCAA) degradation, urea cycle/amino group metabolism, steroid hormone biosynthesis, lysine metabolism, and hormone metabolism in alginate cultured agarose-embedded chondrocytes. A previous study by Zhai *et al* has suggested the ratio of BCAA to histidine as a biomarker for osteoarthritis [35] with altered levels suggesting metabolic stress or disruption in energy homeostasis often seen in OA-affected tissues. Catabolism of BCAA is tightly regulated, and its byproducts feed into the TCA cycle for energy production [36]. There also exists an inter-relationship between the TCA and urea cycle shunt through the intermediary fumarate, a breakdown product of arginosuccinate via arginosuccinate lyase in the urea cycle [37], that feeds into the TCA cycle for energy production. In cellular metabolism, the TCA cycle acts as a central hub providing a source of intermediary metabolites for precursors of biosynthetic pathways, cellular repair, and regeneration.

Interestingly, in monolayer cultured agarose-embedded chondrocytes, urea cycle/amino group metabolism was downregulated along with other pathways such as N-glycan biosynthesis and butanoate metabolism. The downregulation of the urea cycle and amino group metabolism suggests a shift away from protein turnover and biosynthesis. This downregulation of N-glycan biosynthesis could reflect an adaptation in response to altered mechanical signals in the monolayer culture environment. In contrast upregulated pathways were fatty acid metabolism, saturated fatty acid oxidation, and tryptophan metabolism. Fatty acids are a major component of lipids, and their metabolism through beta oxidation is essential for energy regulation [38]. This demonstrates the different energy sources that are used by chondrocytes upon applied cyclical compression when precultured in alginate versus culturing as monolayer.

Chondrocytes precultured in a monolayer appear to adapt by tapping into fatty acid oxidation and tryptophan metabolism as primary energy sources under compression. In contrast, chondrocytes precultured in alginate may retain a more balanced approach to energy generation, incorporating amino acid metabolism and the urea cycle, which could reflect a metabolic profile more aligned with cartilage’s native environment.

Median intensity heat map analysis also found that different pathways were upregulated when chondrocytes were precultured in alginate and then subjected to compression. We observed that when primary chondrocytes were encapsulated in alginate and cultured in the presence of ascorbate, pathways related to amino acids were upregulated.

Of specific importance was the upregulation in the arginine, proline, lysine, and glutamate metabolism. Hydroxyproline and hydroxylysine derived from proline and lysine respectively, are major amino acids found in Collagen VI (Col VI) in addition to glycine. Col VI is a prominent component of the PCM and plays a role in the microenvironment of the chondrocytes. The PCM surrounds the chondrocytes and modulates the biomechanical, biophysical, and biochemical signals [14]. The PCM also directly transduces physical signals from the ECM to the cell, as well as possibly through interactions of Col VI with integrins or other cell surface receptors. It is thought that the PCM plays a role in cell anchoring and matrix cell signaling and is most likely a contact between the hard interterritorial cartilage matrix and the chondrocyte [39].

The observed upregulation of these amino acid pathways is consistent with our immunocytochemistry data that find enhanced production of cell-directed PCM synthesis. Downregulated metabolic pathways for the alginate group were mainly associated with fatty acid oxidation, keratan sulfate biosynthesis, glycan biosynthesis, and metabolism of glycerophospholipids and glycosphingolipids. Lipids are essential components of biological membranes, with glycerophospholipids being the most abundant of the membrane lipids. Both glycosphingolipids and glycerophospholipids play important roles in cellular processes such as cell signaling and energy storage [40]. Interestingly, glycerophospholipid metabolism was upregulated in the alginate-encapsulated chondrocytes, indicating that these membrane phospholipids may differentially modulate signaling cascades when cultured under different conditions.

Keratan sulfate is a widely distributed glycosoaminoglycan present in the ECM of the chondrocytes in articular cartilage and provides compressive resistance to cartilage [41]. Beyond its structural role in cartilage, keratan sulfate also plays a significant role in other tissues. For e.g. in neurons, an additional key role for keratan sulfate lies in sensing ion fluxes allowing them to respond dynamically to biochemical changes within the cell microenvironment [42]. The observed downregulation of keratan sulfate biosynthesis in alginate-cultured chondrocytes may reflect an altered adaptation to the alginate matrix environment, potentially due to reduced mechanical stimuli or differences in nutrient and ion exchange compared to the native cartilage environment.

Metabolism of inflammatory leukotrienes was also down regulated in alginate-encapsulated agarose-embedded chondrocytes. Inflammatory mediators associated with OA often include cytokines, chemokines, prostaglandins, nitric oxide, and leukotrienes [43]. Leukotrienes are lipid-based inflammatory mediators derived from arachidonic acid, and they play a significant role in driving inflammatory responses within tissues. This downregulation suggests a reduced pro-inflammatory profile in these chondrocytes, which may be indicative of a shift in cellular signaling toward a more anabolic or repair-focused metabolic profile. Biosynthesis of coenzyme A (CoA) from its key precursor Pantothenate was also downregulated in the uncompressed alginate encapsulated agarose-embedded chondrocyte group indicating an adaptive response to the lower mechanical demands of the encapsulated environment compared to *in vivo* conditions.

Pathways detected in the monolayer cells embedded in agarose and subjected to cyclical compression included porphyrin, beta-alanine, aspartate and asparagine metabolism, whereas uncompressed pathways were mapped to phytanic acid and peroximal oxidation which is involved in degradation of complex fatty acids. While compression prompts monolayer-cultured, agarose-embedded chondrocytes to upregulate pathways supporting energy production and amino acid processing, uncompressed cells favor pathways involved in lipid metabolism and fatty acid degradation.

Upregulated pathways in the uncompressed alginate cultured-agarose- embedded bovine chondrocytes included amino acid metabolism (lysine degradation, amino acids glycine, threonine, serine and tryptophan), lipid metabolism (sphingolipids), and the phosphatidyl inositol signaling system. Glycine is a primary amino acid found in collagen, and threonine and serine feed into the TCA cycle via pyruvate and acetyl CoA thereby regulating the energy cycle.

Tryptophan metabolism into its downstream metabolites like kyneurenine and derivatives modulate host inflammatory responses by binding to host transcription factor aryl hydrocarbon receptor (AHR) [44]. While AHR is the most described host receptor for tryptophan metabolites, there are other ligand activated transcription factors that play an important role in metabolism of xenobiotics, that can enter animal bodies through feed additives. Since xenobiotic metabolizing enzymes of the cytochrome P450 family are main targets of AHR induction, it validates the upregulation that we see in the metabolism of xenobiotics by cytochrome P450 in bovine chondrocytes [45].

Phosphatidylinositols are a group of phospholipid molecules that are present in cellular membranes and are key regulators of cell signaling and intracellular trafficking [46] whereas sphingolipids are lipid molecules that play critical roles in cell membrane biology to regulate cell function [47]. Energy-related pathways such as starch and sucrose metabolism and ubiquinone biosynthesis were also upregulated in this cluster. Other pathways that were upregulated included starch and sucrose metabolism. Non-structural carbohydrates such as sucrose and starch are metabolized for energy production [48]. Ubiquinone is an electron carrier in oxidative phosphorylation [49] and its biosynthesis was upregulated, contributing to the bioenergetic activity of ATP synthesis. Similarly, the Panthotenate and CoA biosynthesis pathway upregulation in the alginate compressed cluster points to energy production through the TCA cycle. Pathways related to uncompressed monolayer cultured chondrocytes identified primarily as carbohydrate and energy metabolism.

In summary, we present differences in morphology and mechanotransduction pathways when chondrocytes are cultured in monolayer (2D) versus alginate (3D). Chondrocytes cultured in monolayer dedifferentiate and lose their characteristic phenotype over time. However, chondrocytes cultured in alginate maintained their phenotype and produced cartilage-specific PCM components. Similarly, our results indicate that primary human and bovine chondrocytes can form a robust pericellular matrix that mimics the *in vivo* phenotype when cultured in alginate in the presence of sodium L-ascorbate. This dense matrix provides structural support and facilitates the chondrocytes’ response to mechanical stimuli, underscoring the importance of 3D culture systems for studying cartilage biology.

Our study also shows substantial differences in metabolomic profiles of chondrocytes that are encapsulated in alginate compared to monolayer. Many pathways that were detected with the alginate-encapsulated chondrocytes were related to amino acid metabolism, fatty acid metabolism, O and N-glycan biosynthesis, phosphatidylinositol signaling, glycerophopholipid metabolism which is involved in membrane structural integrity and central energy metabolism. These pathways suggest a metabolic profile that supports both ECM synthesis and structural stability in response to the 3D environment of the alginate matrix.

Pathways related to the monolayer cultured agarose-embedded chondrocytes were associated mainly with fatty acid oxidation, terpenoid backbone biosynthesis, porphyrin metabolism (seen also in bovine monolayer cultured agarose-embedded chondrocytes), amino sugar, and monosaccharide (fructose, mannose) metabolism and aspartate and asparagine metabolism. Taken together, imaging data, western blot and amino acid metabolism pathways suggest an increase in collagen production within the pericellular matrix when chondrocytes are cultured in alginate compared to monolayer conditions.

Future studies will investigate the specific pathways that shape chondrocyte biology across these culture systems, offering a clearer understanding of how alginate culture systems support physiological chondrocyte functions. This work not only enhances our knowledge of chondrocyte mechanotransduction but also holds significant potential for cartilage tissue engineering, laying the groundwork for regenerative therapies aimed at cartilage repair and osteoarthritis treatment.

## Acknowledgements

The authors thank Professor Martin Stoddart for helpful discussions that guided this study, which was supported by funding from NIH (NIAMS R01AR073964 and R01AR081489) and the NSF (CMMI 1554708). We thank the Mass Spectrometry Facility at Montana State University in assisting with LC-MS analysis. Funding for the Montana State Mass Spectrometry Facility used in this publication was made possible in part by the MJ Murdock Charitable Trust, the National Institute of General Medical Sciences of the National Institutes of Health under Award Numbers P20GM103474 and S10OD28650, and the MSU Office of Research and Economic Development.

## Author Contributions

PPB designed and executed experiments, analyzed and interpreted data and drafted manuscript. AEE ran the LC-MS and analyzed data. EPM assisted in loading experiment. RKJ designed the experiments, analyzed data, drafted and edited the manuscript. All authors have read and revised the manuscript.

